# Characterizing Cancer Drug Response and Biological Correlates: A Geometric Network Approach

**DOI:** 10.1101/222943

**Authors:** Maryam Pouryahya, Jung Hun Oh, James Mathews, Joseph O. Deasy, Allen Tannenbaum

## Abstract

In the present work, we consider a geometric network approach to study common biological features of anticancer drug response. We use for this purpose the panel of 60 human cell lines (NCI-60) provided by the National Cancer Institute. Our study suggests that utilization of mathematical tools for network-based analysis can provide novel insights into drug response and cancer biology. We adopted a discrete notion of Ricci curvature to measure the robustness of biological networks constructed with a pre-treatment gene expression dataset and coupled the results with the GI50 response of the cell lines to the drugs. The link between network robustness and Ricci curvature was implemented using the theory of optimal mass transport. Our hypothesis behind this idea is that robustness in the biological network contributes to tumor drug resistance, thereby enabling us to predict the effectiveness and sensitivity of drugs in the cell lines. Based on the resulting drug response ranking, we assessed the impact of genes that are likely associated with individual drug response. For important genes identified, we performed a gene ontology enrichment analysis using a curated bioinformatics database which resulted in very plausible biological processes associated with drug response across cell lines and cell types from the biological and literature viewpoint. These results demonstrate the potential of using the mathematical network analysis in assessing drug response and in identifying relevant genomic biomarkers and biological processes for precision medicine.

---

In this paper, we propose the use of certain tools from discrete geometry to gain new insights into cancer drug response. For this purpose, we tested our methodology on the panel of 60 human cancer cell lines (the NCI-60). It has been more than 30 years since the U.S. National Cancer Institute (NCI) established a human cell line panel for the purpose of discovering novel cancer drugs. The NCI-60 panel was designed to recast the previous murine-based drugs from leukemia treatment to the treatment of more diverse human solid tumors. This departure was due to the difference and diversity of the biology of human tumors from murine leukemia (1). This panel was developed as part of the NCI’s Developmental Therapeutics Program (DTP, http://dtp.nci.nih.gov) to screen in *vitro* response to over 100,000 chemical compounds and natural products including FDA-approved anti-cancer drugs and those currently undergoing clinical trial. This ongoing service is accepting global submissions and continues screening up to 3,000 small molecules per year as potential anti-cancer therapies. The 60 human tumor cell lines represent nine tumor types: leukemia, breast, central nervous system, colon, skin, lung, ovarian, prostate, and renal cancers. The NCI-60 panel is thus an established tool for in *vitro* drug screening and has significantly improved the philosophy and research of human cancer drugs (2). This panel has led to many important discoveries, including a general advance in the understanding of the mechanism of cancer and the action of drugs (3, 4). Moreover, comprehensive genomic data including transcript expression data, protein expression data, re-sequencing (mutation) data, DNA copy number, and methylation as well as drug screening GI50 data on the 60 cell lines make it a unique resource for system pharmacogenomics and systems biology (5). Most importantly for our work, this data resource enables us to explore both pre-treatment genomic data and drug responses of a notable number of FDA approved anticancer agents (~130) which is unmatched by any other cancer databases (1).

We are interested in analyzing the gene interaction network for these cell lines via mathematical tools. In recent years, there have been tremendous efforts to elucidate the complex mechanisms of biological networks by investigating the interactions of different genetic and epigenetic factors. Given that gene/protein interactions inherently form a mathematical network, it is reasonable to expect that mathematical tools can facilitate a better understanding of the complexities of such networks (6). The mathematical methods and tools employed in mathematical network analyses are quite diverse and heterogeneous, ranging from graph theory as abstract representations of pairwise interactions to complicated systems of partial differential equations that try to capture all details of biological interactions. The ability to sequence and analyze genomes has revolutionized the diagnosis and treatment of diseases. With the exponential growth of genomic data, the need for improved mathematical methods to analyze the data is becoming even more prevalent. Here, we adopt a discrete mathematical notion of curvature defined on networks to study the robustness of gene interaction networks in response to drugs. We rank the effectiveness of anticancer drugs and find the biological processes in which the important genes are involved. The network based analysis of ‘omics’ allows identification of new disease genes, pathways and rational drug targets that were not easily detectable by isolated gene analysis. This study illustrates the use of a novel mathematical approach to networks to identify pertinent biological processes as well as effective drugs for the treatment of cancer.

The mathematical notions can be summarized as follows. Ricci curvature is a fundamental concept in Riemannian geometry; see (7, 8) for all of the details. Here, we use an analogous notion on discrete spaces, namely, Olivier-Ricci curvature (9, 10). The concept of curvature was initially introduced to express the deviation of a geometric object from being flat. The Riemann curvature tensor of such a manifold encodes key geometric properties and expresses the deviation from Euclidean (flat) space. The sectional curvature is defined on two-dimensional subspaces of the tangent planes, and Ricci curvature is the average of sectional curvatures of all tangent planes containing some given direction (7). Interestingly, Ricci curvature also appears in optimal mass transport theory (11, 12), and serves as the motivation for certain discrete analogues. Indeed, on a Riemannian manifold, one can endow the space of probability densities with a natural Riemannian structure (13, 14) employing the 2-Wasserstein distance from optimal mass transport (15). Thus, given the Riemannian-type metric, one can define a notion of geodesics on the space of probability densities. As noted by Lott-Sturm-Villani (16–18), considering this Riemannian structure, one can relate the Ricci curvature of the underlying manifold, the entropy of densities along a given geodesic path, and the 2-Wasserstein distance in one remarkable formula (see our discussion below for the details). In conjunction with the Fluctuation Theorem (19), we can conclude that increases in the Ricci curvature are positively correlated with increases in the robustness, herein expressed as ΔRic × Δ*R* ≥ 0. Following our previous work (20–22), we are interested in finding important nodes (genes) within the network in terms of robustness.

Coupling the results of the network analysis with the drug growth inhibition values provides us with a network-based guide to the sensitivity/ resistance of the tumor cell lines to these drugs. Here, due to some missing values, we focus on a subset of 58 cell lines. The transcription expression data provided for these cell lines along with the gene-to-gene relationships enables us to construct a weighted network. Investigating this network and relating the information it provides to the drug response gives a novel insight through the NCI-60 database which has not been studied using a network mathematical approach before.

Our main results are based on the application of the aforementioned discrete notion of Ricci curvature to the network generated from the pre-treatment gene expression for all 58 cell lines. This notion allows us to identify possible targets for the anti-cancer drugs. Moreover, for a given drug, we find the average Ricci curvature of the genes whose expressions are significantly correlated to the GI50 response of the drug. In fact, we identify which part of the network is most correlated to a specific drug’s action. The average Ricci curvature for this subnetwork can act as a guide to the sensitivity/resistance of cell lines to the drug. Specifically, the more degree of robustness for the subnetwork can identify the resistance of the drug along the tested cell lines. We are also interested in the biological processes that the significant genes of effective drugs are involved in. This can help us to detect key biological processes associated with the drug response.

The results in the present work are all derived from the network analysis of the NCI-60 genomic information. In our work, we utilized geometric tools in discrete mathematics to better understand these complex networks. This point of view can help to elucidate important drug stratification and biomarkers, which in turn may allow researchers to glean new clinical information from the NCI-60 database.

## Methods

### Background on curvature

In the present study, we employed an analogous notion of Ricci curvature to analyze cancer protein expression networks. In the classical continuous setting, the Ricci curvature tensor provides a way of measuring the degree to which the geometry determined by a given Riemannian metric differs from that of ordinary Euclidean space; see (7) for all the details. We briefly sketch the main ideas to motivate the discrete definition applicable to the networks of interest.

Assume that *M* is a complete connected Riemannian manifold equipped with metric *g*. Positive Ricci curvature at a point of *M* is characterized by the following fact. Let *x, y* ∈ *M* be two very close points defining tangent vector (*xy*). Moreover, let *ω* be a tangent vector at *x* and *ω′* be the tangent vector at *y* obtained by parallel transport of *w* along (*xy*). Then (for positive curvature), the trajectories of the geodesics corresponding to *ω* and *ω′* will approach each other. This can be compared to the traditional flat geometry of a Euclidean space, where such geodesics are always equidistant from each other with their “direction” being unchanged by parallel transport. Equivalently, this may be formulated by the fact that the distance between two small geodesic balls is less than the distance of their centers. Ricci curvature along the direction (*xy*) quantifies this, averaged on all directions *w* at *x*. On the other hand, when the curvature is negative, the geodesics diverge. Lower bounds on the Ricci curvature prevent geodesics from diverging too fast and geodesic balls from growing too quickly in volume (7). In other words, lower Ricci curvature bounds estimate the tendency of geodesics to converge.

Interestingly, optimal transport offers a formulation of lower Ricci curvature bounds in terms of entropy (17, 18). Optimal mass transport theory is concerned with the problem of finding an optimal transport plan (relative to some cost function) for moving a given initial mass distribution (or gene expression levels in our case) *μ* into a final configuration *ν* in a mass preserving manner (11, 12, 16, 23). We will assume that *μ* and *ν* are normalized to be probability measures. Let 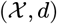 denote a metric measure space. Then the *p*-Wasserstein distance between *μ* and *ν* is defined as

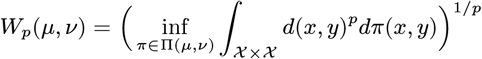

where the latter infimum is taken over all joint probability measures *π* on 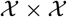 whose marginals are *μ* and *ν*, i.e.:

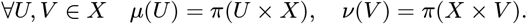

Consider the case 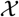 is a Riemannian manifold. The Wasserstein distance defines a Riemannian distance function on the space of the probability measures on 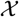 (13, 14). We denote this space by 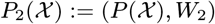. Using the theory of optimal transport, Lott, Sturm and Villani (17, 18) derived an elegant connection between Ricci curvature, Ric, and the Boltzmann entropy, Ent. Namely, Ric ≥ *k* if and only if the entropy functional is displacement *k*-concave along the 2-Wasserstein geodesics, i.e. for all 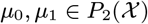 and *t* ∈ [0,1] we have:

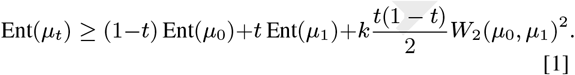

Note that by definition,

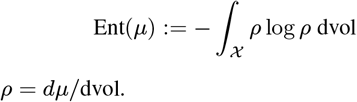

where *ρ* = *dμ*/dvol.

### Robustness defined on networks

The relation (1) indicates a positive correlation between changes in entropy and changes in Ricci curvature that we express as

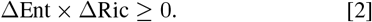

We will describe notions of Ricci curvature and entropy on graphs below. We just note here that changes in *robustness*, i.e., the ability of a system to functionally adapt to changes in the environment (denoted as Δ*R*) is also positively correlated with entropy via the Fluctuation Theorem (19), and thus with network curvature:

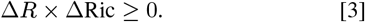

More precisely, the measure of robustness employed in (19) is the *rate function, R*, from the theory of large deviations (24). One considers random perturbations of a given network that result in deviations of some observable. We let *p_ϵ_*(*t*) denote the probability that the mean of the observable deviates by more than *ϵ* from the original (unperturbed) value at time *t*. Since *p_ϵ_*(*t*) → 0, we want to measure its relative rate, that is, we set

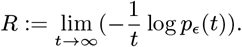

Therefore, large *R* means not much deviation and small *R* large deviation. In thermodynamics, it is well-known that entropy and rate functions from large deviations are very closely related (19). The Fluctuation Theorem is an expression of this fact for networks, and may be written as

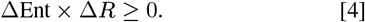

The Fluctuation Theorem has consequences for just about any type of network: biological, communication, social, or neural (19). In rough terms, it means that the ability of a network to maintain its functionality in the face of perturbations (internal or external), can be quantified by the correlation of activities of various elements that comprise the network. In the standard statement, this correlation is given via entropy. This has been reformulated geometrically in terms of curvature (21, 22). We will now give the precise definition for networks modeled as weighted graphs.

### Curvature on weighted graphs

In discrete settings, we assume that our network is represented by an undirected and positively weighted graph, *G* = (*V, E*), where *V* is the set of *n* vertices (nodes) in the network and *E* is the set of edges. We set

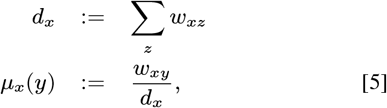

where the sum is taken over all neighbors *z* of *x*, and *w_xy_* denotes the weight of an edge connecting *x* and *y* (it is taken as zero if there is no connecting edge between *x* and *y*).

One of the key characterizations of Ricci curvature on a discrete metric measure space, is via the *Ollivier-Ricci curvature*. As discussed by Ollivier (10) and indicated in Fig. 1, the Ricci curvature of Riemannian manifold can be characterized by comparing the distance between small geodesic spheres and the distance between their centers. Ollivier then extended this idea from the geodesic sphere to an associated probability measure near a point on a metric space 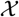.

**Fig. 1.**
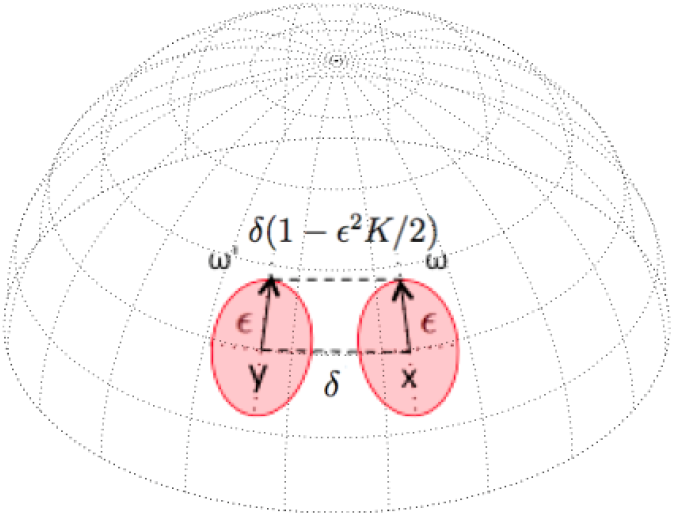
In a positively curved space the distance between the end points of tangent vectors *ω* and *ω′* is less than *δ*. Curvature (*K*) quantifies this difference.

Consider any metric 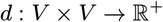 on the set of vertices *V*. For example, *d*(*x, y*) may denote the number of edges in the shortest path connecting *x* and *y*. For any two distinct points *x, y* ∈ *V*, the *Ollivier-Ricci (OR) curvature* is defined as follows:

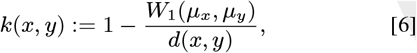

where *μ_x_, μ_y_* are defined in (5). In the present study, we used the Hungarian algorithm (25) to compute the Earth Mover Distance on our reference networks. This discrete notion of Ricci curvature has been already used to investigate the robustness of cancer networks (21, 22).

Using this edge based notion of curvature, we can also define the *scalar curvature* of a given node in the graph as follows:

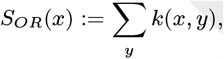

where the sum is taken over all neighbors of *x*.

### Gene expression and drug activity data

The mRNA expression data for the NCI-60 human tumor cell lines were retrieved from the CellMiner (http://discover.nci.nih.gov/cellminer). Cellminer is a web application written by the Genomics & Bioinformatics Group, LMP, CCR, NCI (5), which provides freely accessible analysis tools and downloadable data sets for exploring NCI-60 data. The database contains transcript expression values for several assays of the NCI-60 cell lines. This study utilizes Affymetrix HG-U133 (A-B) with GeneChip RMA (GC-RMA) normalization from this website.

Using these gene expressions arrays, we found the gene-to-gene correlations to build our weighted networks. The underlying topology has been derived from Human Protein Reference Database (HPRD, http://www.hprd.org) (26). Affymetrix HG-U133 (A-B) data contains 34,899 probes. For the repeated gene IDs, we found the average RNA expression of the corresponding probes. We used only probes with known gene names. Resulting in 16,821 genes with RNA expression. We chose the intersection of these genes with the HPRD database as the nodes of the pre-treatment network. Overall, the network consists of 8240 genes. The weights of the edges are defined by the Pearson correlation between the gene expressions along the 58 cell lines. We further used the transformation of 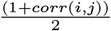 for the genes *i* and *j* to make a positively weighted network. We then found the Ollivier-Ricci curvature for all the edges, and the scalar curvature for all the nodes (genes) within the pre-treatment network. We further identify significant genes (nodes) for each drug in this network. These important genes were selected based on significant Spearman’s correlations (*p*-value < 0.05) between drugs’ activity, 50% growth inhibition (GI50), and gene expressions across the cell lines. Finally, we computed the average of Ricci curvature values for the significant genes associated with each drug.

The GI50 is the drug concentration resulting in a 50% reduction in the net protein increase during drug incubation as compared with the same increase in control cells (27, 28). Normalized (− log_10_) GI50 was retrieved using R package, *rcellminer* for exploring CellMiner database. This package complements the functionality of CellMiner by providing programmatic data analysis and visualization (31). Fig. 3(b) derived from the rcellminer package illustrates the Spearman’s correlation (Rs) between the gene expression of MYC (z-score) and methotrexate activity (z-score). In this case, the correlation is very significant (*Rs* = 0.41) with a very low *p*-value (0.001). Finding the correlation between all selected genes and FDA approved drugs, we constructed the Spearman’s correlation matrix shown in Fig. 2. There were no expression information of Affymetrix HG-U133 (A-B) for LC:NCI-H23 in non-small cell lung cancer (NSCLC), and many missing GI50 values for the drug responses of ME:MDA-N of the melanoma cell line. Excluding these two cell lines resulted in 58 complete data sets (see Fig. 3(a)).

**Fig. 2.**
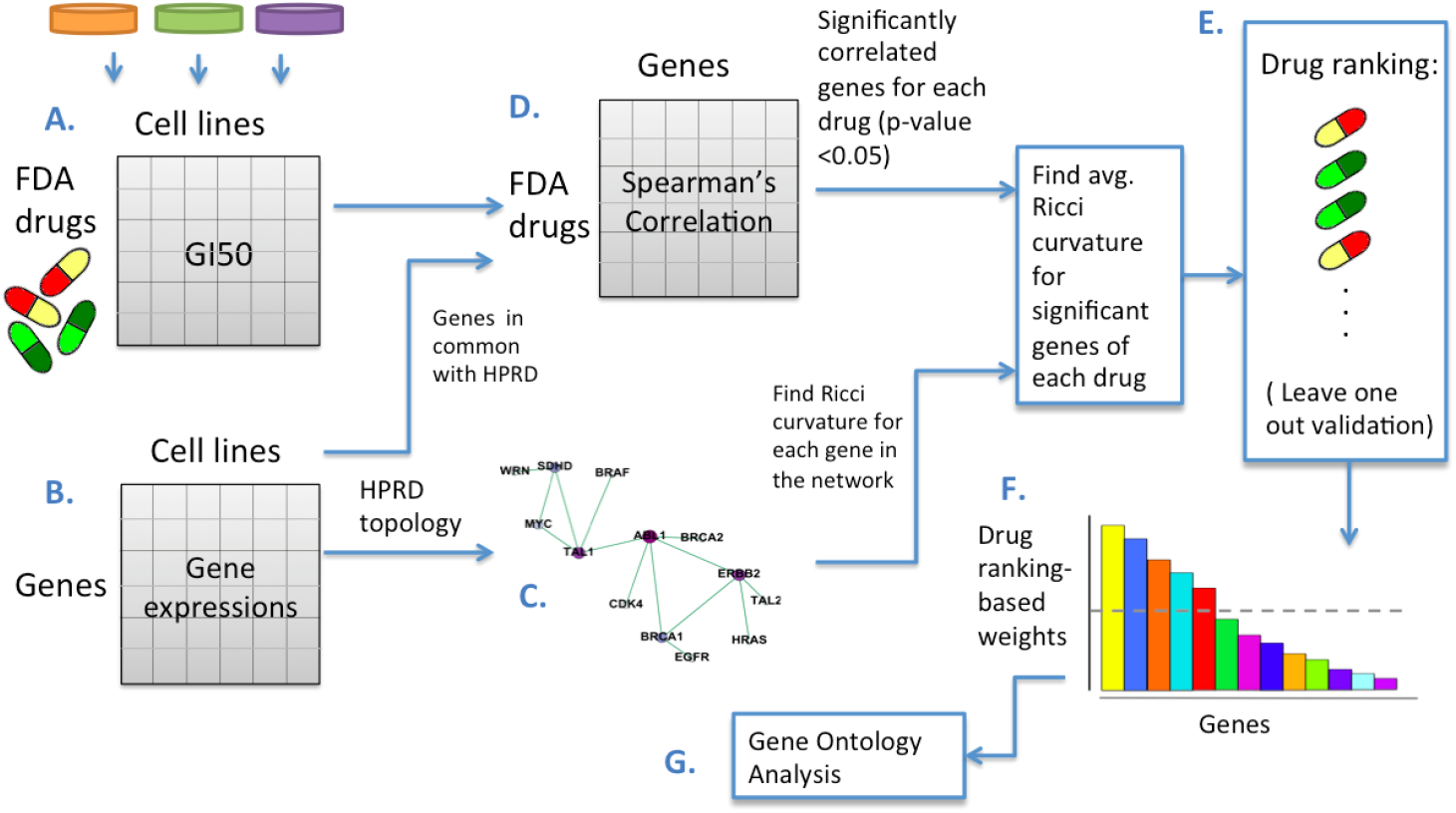
Methodology for establishing a network-robustness ranking of genes across cancer drugs and cancer cell lines: A. GI50 drug activity matrix of 129 drugs for 58 cell lines. B. Matrix of 8240 gene expressions for 58 cell lines. C. Pre-treatment network made by the gene expression correlation along 58 cell lines as the weights, and the underlying topology of gene-to-gene interactions is derived from HPRD. D. Matrix of Spearman’s correlations between each drug’s activity (rows of matrix A.) and gene expression (rows of matrix B.) along 58 cell lines. E. Ranking drugs in ascending order of average Ricci curvature values of significant genes. F. Drug ranking used to score the significant genes correlated to the drugs. G. Top 200 genes selected for gene ontology enrichment analysis.

**Fig. 3.**
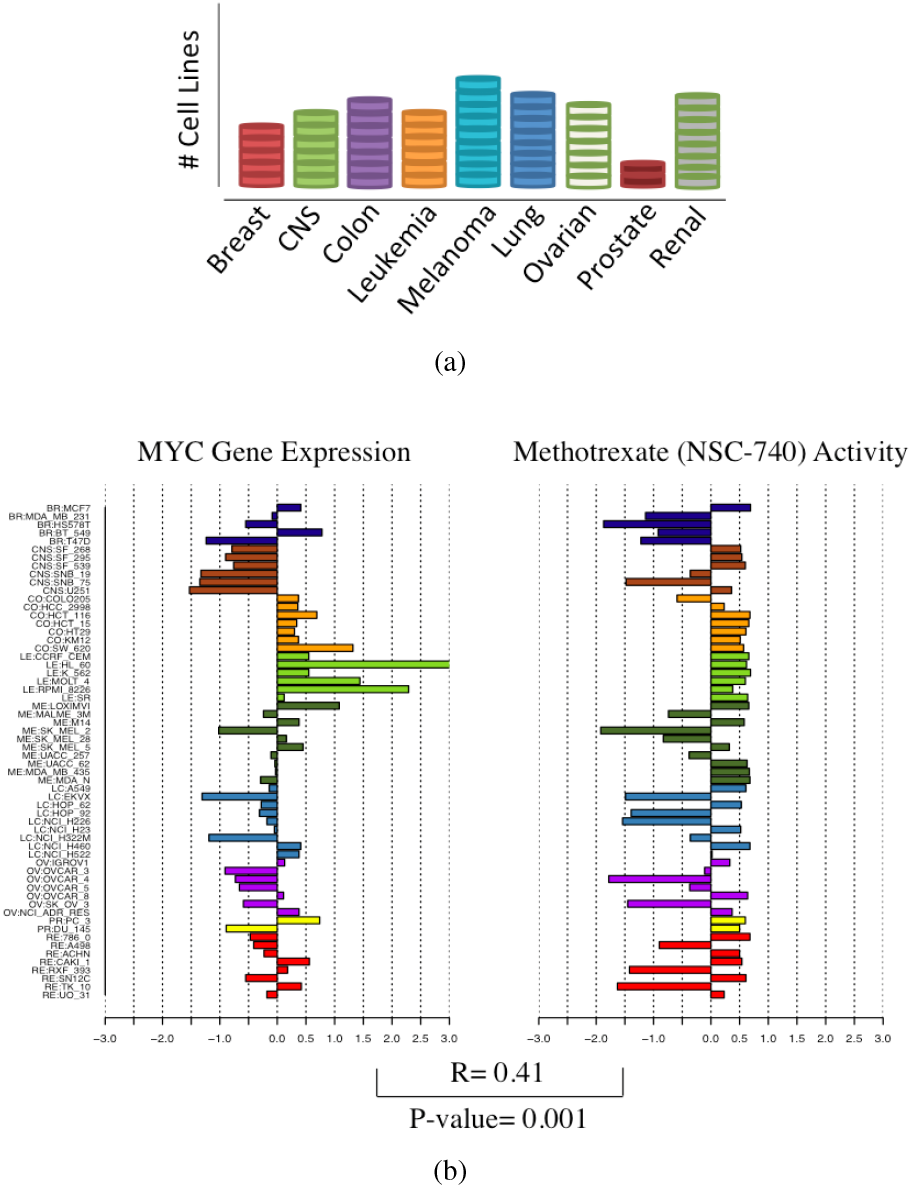
(a) 58 cell lines distribution (b) Using expression data such as this example (Methotrexate and MYC), Spearman’s correlation between each drug activity’s GI50 and gene expression along cell lines has been calculated to create the Spearman’s correlation matrix (D) shown in Fig. 2.

### Ranking drugs and gene ontology enrichment analysis

The summary of our methodology is shown in Fig. 2. The FDA approved drugs correspond to 161 NSC drug numbers (numeric identifiers for substances submitted to the National Cancer Institute) with less than 2 missing GI50 values and 129 drugs with no missing values across all the cell lines. For each drug, we identified genes whose expressions were significantly correlated (*p*-value < 0.05) to GI50. The median number of the genes selected for each drug was ~500 with the minimum number ~300. The average Ollivier Ricci curvature of these genes was calculated for each drug.

We then sorted the 129 FDA approved drugs in ascending order of the average Ricci curvature (Fig. 2E) arguing that the sensitivity of drugs in cell lines is positively correlated with the changes in robustness of the nodes it perturbs in the network. For the repeated drugs, the GI50 data were different for the different NSC numbers, and therefore, they have different rankings. Although the rankings were close for these drugs, we generally chose the highest ranking as a representative for the repeated drug. Consequently, our final ranking consists of 106 drugs. We primarily performed most of the programming via Matlab.

We used the drug ranking to assess the importance of 8240 genes in our network across cell lines. More precisely, we gave a linear weight of (129 − r) + 1 to all the selected genes associated with the r-th ranked drug. Then, for each gene, we computed the sum of all these weights and ranked the genes in descending order of the total weights. Thus, the final gene score increases if a gene (a) is important for many drugs, and (b) strongly contributing to robustness across multiple drugs. For a biological analysis, the top 200 genes were selected where the histogram has an apparent sharp decline; see Fig. 5. We performed a gene ontology (GO) enrichment analysis on the top 200 ranked genes using the MetaCore software (Thomson Reuters). The MetaCore is an integrated software based on a manually-curated database of molecular interactions, molecular pathways, gene-disease associations, chemical metabolism and toxicity information.

## Results

We found the scalar Ollivier-Ricci curvature of all the 8240 genes in the pre-treatment interaction network discussed previously. The scalar curvatures range between −210.4 and 3.6 with an average of −5.2. Table S1 presents the top 40 genes with the highest absolute value of Ricci curvature. The top two genes, TP53 and YWHAG, stand out with regards to their Ricci curvatures (See Fig. S1). A visualization of the pre-treatment network is provided in Fig. S2. We then rank the drugs based on the average Ricci curvature of significant genes for each drug. The top 30 ranked drugs are presented in Table 1. There are a number of very effective drugs that are ranked highly in this table; we elaborate on these drugs further in the discussion section below. The ranking of all 106 drugs has been presented in Table S3.

**Table 1.**
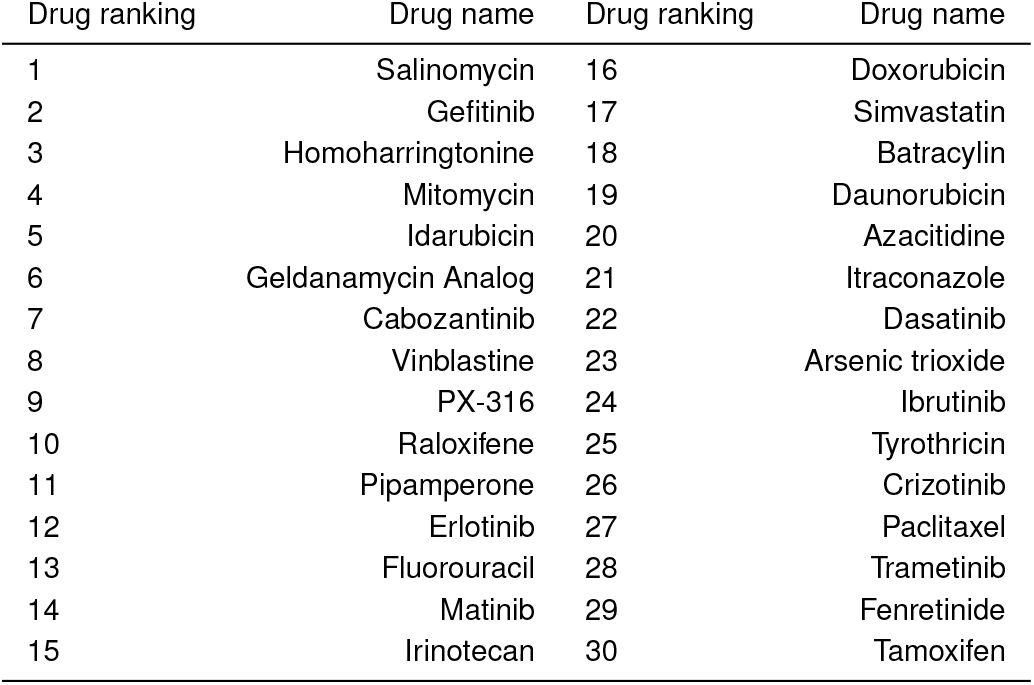
Top 30 drugs ranked by average Ricci curvature of significantly correlated genes.

We validated this ranking by running the entire algorithmic pipeline nine times; see Fig. 2. Each time we excluded all the cell lines of one of the nine cancer tissues. We then computed the Spearman’s correlation between drug rankings resulting from all the cell lines and those based on leave-one-out rankings. Interestingly, correlations are very high with low p-values, showing relative consistency of the results across cell lines. The color map in Fig. 4(b) illustrates these correlations. The rows (columns) of this symmetric map are numbered by excluding all the cell lines of the corresponding cancer tissue shown in the pie chart in Fig. 4(a). For example, w1 corresponds to exclusion of all the five cell lines of breast cancer. The 10th row (column) corresponds to the case of considering all the cell lines. The correlation values are also included in the color map. As we see in Fig. 4, the correlation is very significant among the rankings (*p*-value << 0.05), yet it is less significant after excluding the cell lines of leukemia (4th column or row). In other words, ranking of the drugs after exclusion of cell lines of leukemia is different from excluding any of the other eight caner cell lines. This highlights that the effect of leukemia in the drug ranking is different than other cancer tissues. Of note, this is not a computational effect alone, given that leukemia is not even the cancer type with the greatest number of cell lines (colon, melanoma, lung and renal have more cell lines; see Fig. 3(a)).

**Fig. 4.**
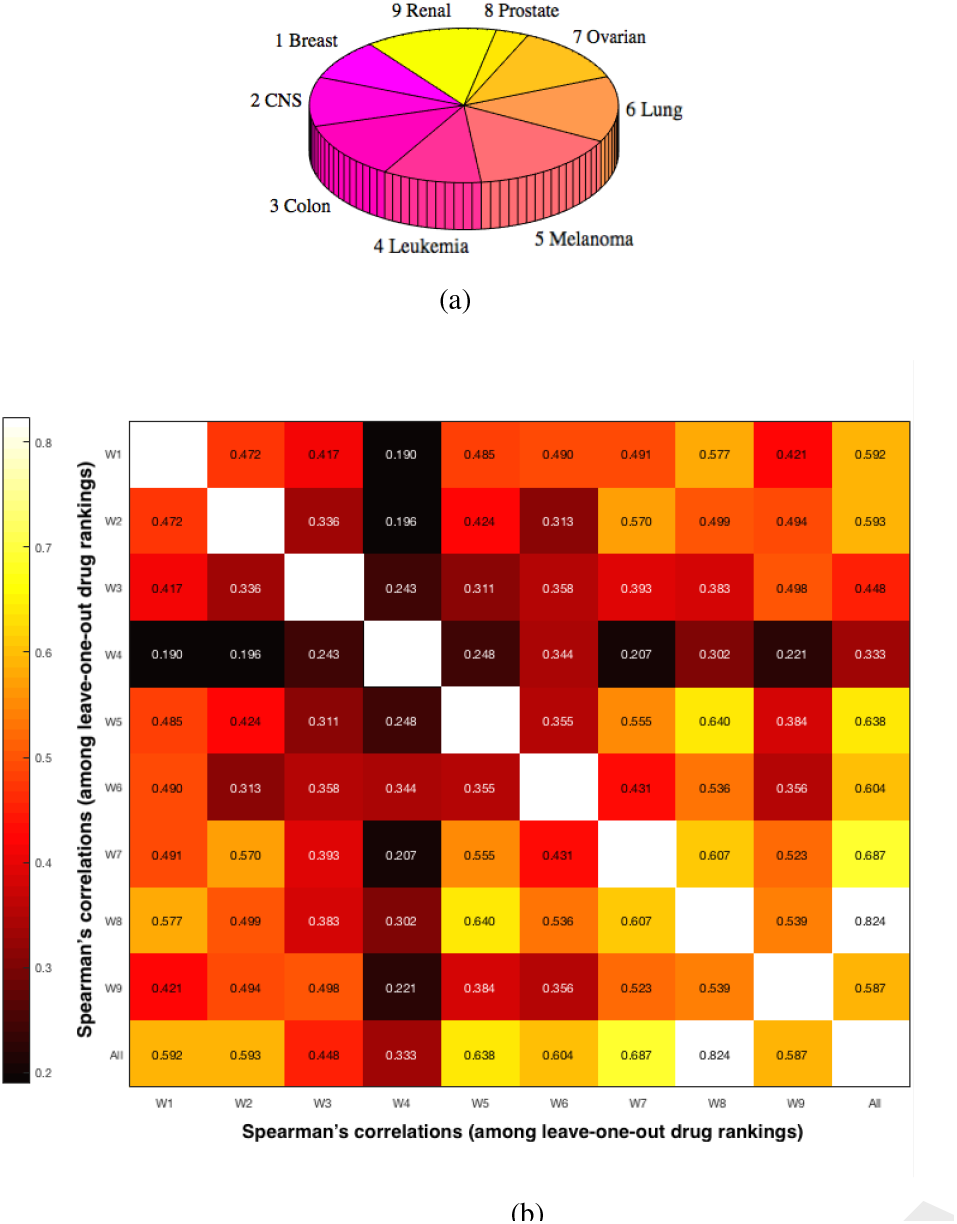
Color map numbered by excluding all the cell lines of the corresponding cancer tissue in the pie chart, (a) Pie chart of cell lines distribution (b) Spearman’s correlation is high between drug rankings.

The top 200 ranked genes, which were selected based on the drug ranking, are presented in Table S3. The results of the gene ontology enrichment analysis of this 200 gene set are shown in Fig. 6(a). The top ten biological processes presented with very small *p*-values. The top three biological processes are all involved with cellular localization. Also, the protein-protein interaction network of this analysis is presented in Figure 6(b). The network contains two hubs associated with the gene product of CUX1 and PRKACA which will be elaborated upon in the discussion section. In addition to analysis of all cancer cell lines, we performed our algorithmic pipeline for three specific cancer tissues with the greatest number of cell lines after melanoma: colon (7 cell lines), lung and renal (both 8 cell lines) cancers. We performed the gene ontology enrichment analysis using the top 200 genes (Table S3) for the cell lines of these specific cancer tissues, and compared the results to the biological processes of all the cell lines. We present the results of the gene ontology enrichment analysis of the top 200 genes of colon, lung and renal cancer tumors in Figure S3, S4, and S5. Interestingly, these cancer types share some similar biological processes to those resulting from all 58 cell lines, which we will discuss further in the next section.

## Discussion

In the present study, we considered the NCI-60 panel comprised of 58 individual cancer cell lines derived from nine different tissues (breast, brain, colon, blood, skin, lung, ovarian, prostate and kidney). The gene expressions of the cell lines were used to construct the network consisting of 8240 genes. This is a weighted graph where the underlying interactions are derived from Human Protein Reference Database. The weights are the gene-to-gene (Pearson) correlations. The network was analyzed by calculating the discrete Ricci curvature. Based on our arguments in the Methods section, we claim that this could be a guide for the robustness of the network, i.e., the ability to withstand perturbations in the system. In our case, these perturbations are induced by the drugs.

The scalar Ricci curvature helps us to identify the important targets (genes) for the drugs within the network. We present the top 40 genes with the least negative (highest absolute value) of Ricci curvature in the pre-treatment network in table S1. The top two genes, TP53 and YWHAG have very low scalar Ricci curvatures compared to other genes (See Table S1). Of note, mutations of TP53 are present in more than 50% of human cancers, making it the most common genetic event in human cancer (29, 30). This gene has many connections to other genes within the network which also contributes to its extreme value of scalar Ricci curvature (See Figure S2). Even though our primary focus in this study is to identify the important genes based on drug response (Figure 5) and the biological processes they are involved in (Figure 6), we briefly discuss the roles of TP53 and YWHAG (top 2 ranked genes in the pre-treatment network) in cancer pathogenesis in the supporting information section.

**Fig. 5.**
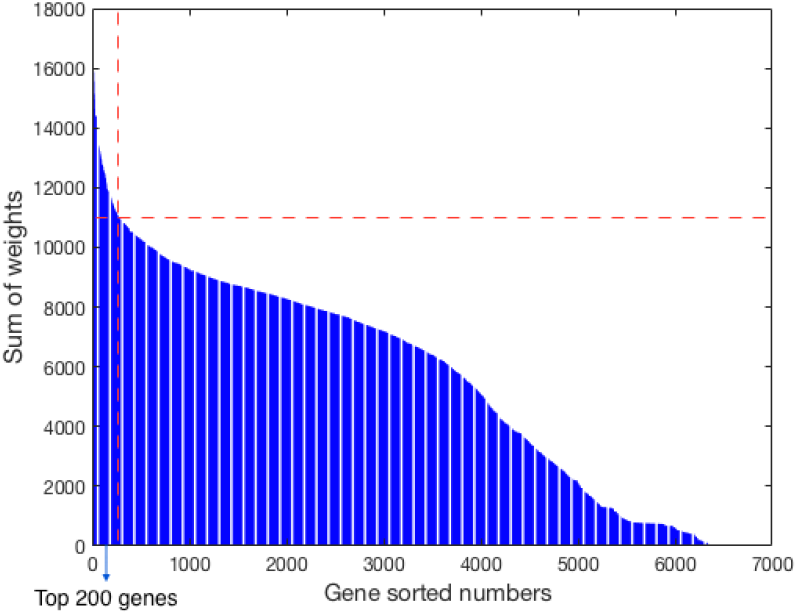
Top 200 genes selected for the gene ontology enrichment analysis.

**Fig. 6.**
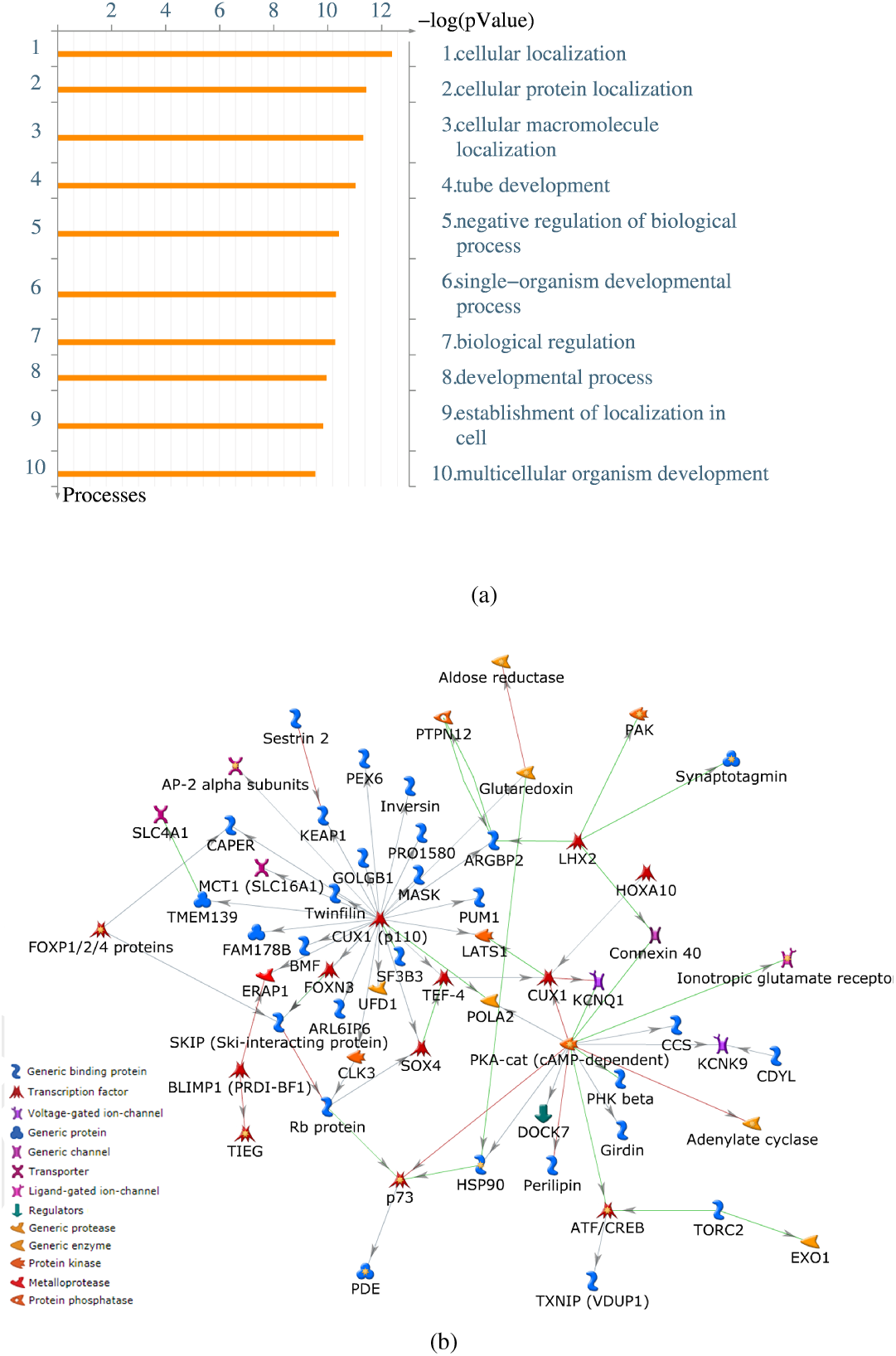
Gene ontology enrichment analysis (MetaCore) of the significant genes correlated with the top ranked drugs (a) Top ten biological processes; top three biological processes are involved with the cellular localization. (b) Protein-protein interaction network has two hubs: CUX1 and cAMP-dependent protein kinase.

Furthermore, we measure the effectiveness of the drugs by the average Ricci curvature of the nodes (genes) it affects. For each drug, we found the significantly (positive/ negative) correlated genes by computing the correlation between GI50 and gene expression along the 58 cell lines. The significant genes for each drug were chosen based on *p*-values less than 0.05. On average ~500 genes were selected for each drug. The average discrete Ricci curvature of these genes was calculated for each drug. Since Ricci curvature and robustness are positively correlated, we were able to rank the network robustness of the drugs by calculating the average Ricci curvature. When the network has higher curvature, we expect that it should show more resistance to the drug. Therefore, the ranking of the drugs in ascending order (of average Ricci curvature), can guide us to the efficacy of the drugs for the cell lines. This network based view of the drug’s effectiveness considers the gene interactions of the cell lines as well as the drug response. The table of top 30 drugs is presented in Table 1. We also provide the table of all 106 drug ranking in supporting information, Table S3.

Salinomycin, the first ranked drug, has recently been considered as a promising novel anti-cancer agent for targeting human cancer stem cells despite its not well-known mechanism of action (32–34). The chemotherapeutic property of Salinomycin can overcome the resistance of tumor cells toward multiple drugs while selectively targeting the cancer stem cells. This antibiotic drug has been shown to kill breast cancer stem cells in mice at least 100 times more effectively than the known anti-cancer drug Paclitaxel. The study screened 16,000 different chemical compounds and found that only a few drugs, including Salinomycin, targeted cancer stem cells responsible for metastasis (35).

Gefitinib, our second ranked drug, is a molecular targeted drug in the treatment of non-small cell lung cancer. Approximately 85-90% of lung cancer cases, the most deadly cancer in the US, are NSCLC tumors. Mutations in the EGFR (epidermal growth factor receptor) gene are present in about 10 percent of NSCLC tumors (https://www.iressa-usa.com). EGFR overexpression leads to inappropriate activation of the anti-apoptotic Ras signalling cascade, thereby leading to uncontrolled cell proliferation (36). Gefitinib competes with adenosine triphosphate at the ATP binding site in epithelial cells, blocking its tyrosine kinase activity, and consequently inhibiting EGFR signaling pathway, which can induce tumor cell apoptosis (37). In 2015, gefitinib was FDA approved as a first line treatment in patients with metastatic NSCLC who harbor the most common types of EGFR mutations in NSCLC (exon 19 deletions or exon 21 L858R substitution gene mutations) (38).

Omacetaxine mepesuccinate, also known as homoharringtonine, the 3rd ranked drug, was originally identified over 35 years ago as a novel plant alkaloid with antitumor properties. Its mechanism of action is thought to be inhibition of protein translation by preventing the initial elongation step of protein synthesis via an interaction with the ribosomal A-site (39). It was approved by the FDA in October 2012 for the treatment of adult patients with chronic myeloid leukemia (CML) with resistance and/or intolerance to two or more tyroskine kinase inhibitors, the current first-line treatment (40). Furthermore, clinical studies have shown activity of omacetaxine in other malignancies as a single agent or in combination with other therapies in acute myeloid leukemia (AML) and myelodysplastic syndrome (MDS), and studies are ongoing in this regard (41–43). Also, a number common antitumor agents were ranked highly: doxorubicin, paclitaxel, fluorouracil (5-FU) and tamoxifen are commonly used in breast cancer treatment. Paclitaxel, vinblastine and irinotecan are often used in NSCLC. Homoharringtonine, azacitidine, and arsenic trioxide are common anticancer agents against leukemia.

The leave-one-out validation of drug rankings suggests that the orders are not highly dependent on specific cancer tissue. As in clinical practice, this supports the use of anticancer drugs for the treatment of different cancer types. Overall, the Spearman’s correlations are higher among solid tumors as compared to the liquid tumor, Leukemia. Network-based analysis can also help to understand important biological processes in which the most correlated genes of the top ranked drugs are involved. To this end, we ranked the genes by using our drug ranking. The genes that are correlated to the top ranked drugs are given higher weights. The genes were then ranked in the descending order of the sum of the weighted values. Therefore, the top ranked genes are more correlated with the drug sensitivity in the cell lines and are better targets in the network. The gene ontology analysis, performed via MetaCore, was employed to find the biological processes with which the top 200 genes are involved.

The top three biological processes are concerned with cellular localization. Cells consists of many different compartments that are specialized to carry out various tasks. Based on the gene ontology directory, cellular localization of a protein is involved with the process whereby a protein complex is transported to, and/or maintained in, a specific location within a cell, including the localization of substances or cellular entities to the cell membrane. The number of proteins that have reliable subcellular location annotations is approximately 20% of all known proteins to date (44). It is of particular interest to determine if a potential target is a cell surface or secreted molecule which would be more easily accessible for the targeted drug approach (45). Therefore, knowledge of the subcellular localization of a protein can significantly improve target identification during the drug development process (46). In our framework, these genes are significant for the top ranked drugs since they were ranked based on the robustness of the drug networks. In other words, since our ranking of the drugs is in the ascending order of average Ricci curvature values of significant genes, the top drugs are involved with the genes that contribute to the network robustness and are likely to be targeted by the drugs.

Furthermore, the protein-protein interaction network of the gene products is presented in Figure 6(b). There are two notable hubs in this network: CUX1 and cAMP-dependent protein kinase which is a gene product of PRKACA. CUX1 is specifically important since it is a transcription factor to a number of our top ranked genes associated with the drug sensitivity. CUX1 (also known as CUTL1) is a homeobox transcription factor highly evolutionarily conserved and plays a known role in embryonic development, cell growth and differentiation in mammals (48). Moreover, the role of CUX1 in drug resistance/sensitivity, has been explored. Specifically, gain-of-function as well as loss-of-function studies have shown that increased CUX1 activity significantly enhanced cell sensitivity and cancer tissue response to chemotherapy drugs and resulted in increased apoptosis and growth inhibition. In contrast, decreased CUX1 expression reduced cell sensitivity to chemotherapy drugs with fewer apoptoses and resultant drug resistance (49). These studies, which have been in the context of gastric cancer, suggest an inverse association between CUX1 and drug resistance, implying that CUX1 is an attractive therapeutic target. Whether this phenomenon applies to cancers other than gastric cancer remains to be elucidated. The human PRKACA gene encodes the PKA catalytic subunit alpha (C_*α*_) isoform. With regards to drug sensitivity/resistance, PRKACA is over expressed in invasive and anti-HER2 therapy (trastuzumab/ lapatinib)-resistant breast cancers. In addition to PRKACA conferring resistance to anti-HER2 therapy, it also impairs apoptosis (50). Consequently, inhibition of PRKACA and/or its downstream anti-apoptotic effectors in combination with anti-HER2 therapy may increase the drug sensitivity. Its role in drug sensitivity/resistance for other cancers needs to be studied further.

Finally, we were interested in comparing the results of the gene ontology enrichment analysis of all 58 cancer type cell lines to some specific cancer tissues. The repeated algorithmic process for colon, lung and renal cancers yields consistency. Similar to all cancer type’s genes, the gene ontology enrichment analysis for the top genes involved with renal and lung cancers results in cellular localization and cellular component organization. However, most of the biological processes from the colon cancer gene ontology enrichment analysis are quite general such as negative regulation of cellular process, which is also a top biological process of all cancer types as well as renal cancer. Of note, colon cancer has fewer cell lines than lung and renal cancer.

The need for network based techniques is becoming more and more prevalent as a result of the exponential growth of data in the genomic era. In this work, we utilized mathematical techniques to drive a network based analysis in order to explore the genomic and pharmacogenomics information of NCI-60. The framework of this study can be extended to find possible optimal combinations of drugs. Combining anti-cancer agents, whether cytotoxic or molecularly targeted, with different mechanisms of action is the most practical approach to overcome single drug resistance and produce sustained clinical remissions. Also, the study described in the present work may be extended to tissue-specific drugs employing a more appropriate tissue-specific cell line database. This analysis on the NCI-60 is promising and supports efforts to analyze larger datasets with advanced network mathematics.

## Acknowledgements

This project was supported by AFOSR grants (FA9550-15-1-0045 and FA9550-17-1-0435), ARO grant (W911NF-17-1-049), grants from the National Center for Research Resources (P41-RR-013218) and the National Institute of Biomedical Imaging and Bioengineering (P41-EB-015902), NCI grant (1U24CA18092401A1), NIA grant (R01 AG053991), MSK Cancer Center Support Grant/Core Grant (P30 CA008748), and a grant from the Breast Cancer Research Foundation.

